# A dynamic network model can explain temporal receptive fields in primary auditory cortex

**DOI:** 10.1101/465047

**Authors:** Monzilur Rahman, Ben D. B. Willmore, Andrew J. King, Nicol S. Harper

## Abstract

Auditory neurons encode stimulus history, which is often modelled using a span of time-delays in a spectro-temporal receptive field (STRF). We propose an alternative model for the encoding of stimulus history, which we apply to extracellular recordings of neurons in the primary auditory cortex of anaesthetized ferrets. For a linear-non-linear STRF model (LN model) to achieve a high level of performance in predicting single unit neural responses to natural sounds in the primary auditory cortex, we found that it is necessary to include time delays going back at least 200 ms in the past. This is an unrealistic time span for biological delay lines. We therefore asked how much of this dependence on stimulus history can instead be explained by dynamical aspects of neurons. We constructed a neural-network model whose output is the weighted sum of units whose responses are determined by a dynamic firing-rate equation. The dynamic aspect performs low-pass filtering on each unit’s response, providing an exponentially decaying memory whose time constant is individual to each unit. We find that this dynamic network (DNet) model, when fitted to the neural data using STRFs of only 25 ms duration, can achieve prediction performance on a held-out dataset comparable to the best performing LN model with STRFs of 200 ms duration. These findings suggest that integration due to the membrane time constants or other exponentially-decaying memory processes may underlie linear temporal receptive fields of neurons beyond 25 ms.

**AUTHOR SUMMARY:** The responses of neurons in the primary auditory cortex depend on the recent history of sounds over seconds or less. Typically, this dependence on the past has been modelled by applying a wide span of time delays to the input, although this is likely to be biologically unrealistic. Real neurons integrate the history of their activity due to the dynamical properties of their cell membranes and other components. We show that a network with a realistically narrow span of delays and with units having dynamic characteristics like those found in neurons, succinctly models neural responses recorded from ferret primary auditory cortex. Because these integrative properties are widespread, our dynamic network provides a basis for modelling responses in other neural systems.

## INTRODUCTION

The response properties of auditory neurons is commonly described by a spectro-temporal receptive field (STRF) model, which characterizes the linear dependence of the neural response on the sound spectrum at a range of latencies (1–14). A static non-linearity is often applied to the output from the linear STRF - this linear non-linear (LN) model estimates the response properties of neurons significantly better than the linear estimate (15,16). While STRF models are somewhat successful in explaining the dependence of neural responses on the past few hundred milliseconds of stimulus history (17), these models do not show how this temporal aspect of receptive fields might be implemented biologically. One naïve possibility is that auditory cortical neurons receive inputs at a range of simple delays spanning out to 200 ms, a direct analogue of STRF models. However, the onset latencies of neurons in the ventral division of the medial geniculate body (18), which provides the primary ascending input to primary auditory cortex (A1) are typically less than 30 ms. This is far less than the duration of stimulus history which influences the responses of A1 neurons.

In this study, we asked how much of the dependence on stimulus history (temporal receptive fields) of neurons in primary auditory cortex can instead be explained by certain simple dynamical aspects of neurons – an approach more consistent with the known biology. To model a neuron’s response, we constructed a neural network model whose output is the weighted sum of the responses of multiple units, each of which resembles an LN model. However, the response of each unit of the network was modified in accordance with a dynamic firing-rate equation. This low-pass filters the unit’s response (by convolving with an exponential decay impulse response), providing a simple exponentially decaying memory. This integrative characteristic can be related to the capacitance and resistance of nerve cell membranes, and the individual time constant of each unit can be related to the membrane time constant of real neurons (19). The dynamic aspect of our network can alternatively be interpreted in terms of other neurobiological dynamical phenomena, such as channel-based neural adaptation, short term synaptic plasticity, or recurrent network properties. We found that our biologically-motivated dynamical model can accurately capture the temporal receptive fields of neurons without the need for a wide span of latencies of input.

## MATERIALS AND METHODS

### Stimuli

Models were fitted to single-unit neural responses of anesthetized ferrets to natural sound stimuli. Altogether, 20 sound clips were presented containing human speech in different languages, other animal vocalizations (e.g. ferret) and environmental sounds (e.g. wind and water). All clips were 5 s long with a sampling rate of 48,828.125 Hz and with root mean square intensity ranging from 75 to 82 dB SPL.

### Experimental setup and neural responses

The electrophysiological data used in this study were taken from a series of experiments used in a previous study by Harper et al. (20). Briefly, recordings were made from the primary auditory cortical areas, A1 and the anterior auditory field (AAF) of 6 adult pigmented ferrets (5 females and 1 male) under ketamine (5 mg/kg/h) and medetomidine (0.022 mg/kg/h) anaesthesia for 20 repeats of the 20 natural sound clips played in random order. All animal procedures were performed under license from the United Kingdom Home Office and were approved by the local ethical review committee. In total, 56 penetrations resulted in 549 single and multi-units, of which 284 were single units. A single unit was taken for analysis only if its activity was driven by the stimulus according to the noise ratio (see below). For each unit, the number of spikes was counted in each 5 ms time bin and averaged over repeats, to provide a response profile *y_n_*(*t*), where *t* is time, and *n* is the clip number. The total number of time bins in a clip is *T*. For simpler notation, we will drop the subscript *n*, unless we note otherwise.

### Noise ratio

The noise ratio was calculated as a measure of how much the response of each unit is dependent on the stimuli (9,21). The noise ratio was measured over all 20 stimuli and 20 repeats. Any unit with a noise ratio > 40 were excluded from the study. Also, only putative single units were used. This resulted in 73 neurons for the study.

### Cochleagrams

The models receive the input as a cochleagram, a spectrogram-like transformation of the sound waveform. The cochleagram approximates the spectral filtering performed by the auditory periphery and was calculated as follows (20,22). For each sound clip, the amplitude spectrum was measured using 10-ms Hanning windows, overlapping by 5ms. The number of frequency channels was then reduced by weighted summation using overlapping triangular windows to provide 34 log-spaced channels (500 Hz to 22,627 Hz center frequencies, adapted from melbank.m (http://www.ee.ic.ac.uk/hp/staff/dmb/voicebox/voicebox.html)). Next, a log function was applied to the value in each time-frequency bin, and any values below a low threshold was set to the threshold. The cochleagram was normalized to zero mean and unit variance over the training set (see Cross-validation and model testing). The input *x_fq_*(*t*) at time *t* was the chunk of cochleagram preceding *t*, given by *x_fq_*(*t*) = *K_f_*(*t* – *q* + 1), where *f* is the frequency channel and *q* is the latency and *K_f_*(*t*) is the cochleagram.

### The LN model

The two stage LN model consists of a linear model followed by a sigmoid output nonlinearity.

*Linear stage:*

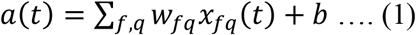

where *a(t)* is the model neuron’s activation, *w_fq_* is the input weight and *b* is the baseline activation. Both *w* and *b* are free parameters of the model and were estimated by regressing *y(t)* against *x(t)* using glmnet (23). To overcome overfitting, the weights were regularized using an L1-norm (LASSO regularization). Thus *a(t)* can be seen as the best linear estimate of *y(t)* from *x(t)*. The regularization hyperparameter *λ* controlled the strength of the regularization. To select *λ*, the model was subjected to k-fold crossvalidation (see Cross-validation and model testing).

*Nonlinear stage:* The nonlinear stage of the model was a parameterized sigmoid nonlinearity:

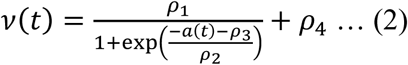

The four parameters *ρ_i_* of the function were fitted by minimizing the squared error between *v(t)* and *y(t)*.

### The Network Receptive Field (NRF) model

The NRF model is an artificial neural network with a single hidden layer of 20 hidden units (HU) which converge onto an output unit (OU). Each unit of the model is similar to a single LN model. The activation of the *j*-th HU, *a_j_*(*t*) is,

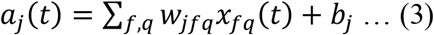

The output of the *j*-th HU, *v_j_*(*t*) is,

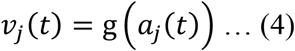

where g (*a_j_*(*t*)) is a sigmoid non-linear activation function, 1/(1 + exp(– *a_j_*(*t*))). The HUs feed these outputs to the OU. The activation function of the OU *a_o_*(*t*) is,

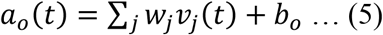

The output *v_o_*(*t*) of the OU is,

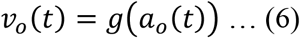

which is the model’s estimate of the neuron’s response at time *t*.

### The Dynamic Network (DNet) model

A neuron’s response depends on the activity of its presynaptic neurons through the total synaptic current they evoke. However, a neuron’s membrane potential, and consequently its firing rate, is not an instantaneous function of this current. Due to the capacitance and resistance of the neural cell membrane, the membrane potential and hence the firing rate is approximately a low-pass filtered version of the synaptic current. A common simple dynamic model of neurons that captures this phenomenon is the differential equation (Equation 7) known as the firing-rate equation (19).

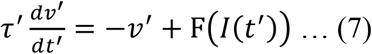

Here, *τ′* is the membrane time constant, *v′* is post-synaptic firing rate, *I*(*t′*) is the synaptic current at time *t′*, F() is a nonlinear function that relates synaptic current to the post-synaptic firing rate, and *t′* is time. Here we use *t′*, *τ′* and *v′* with a prime to denote that for this equation these variables operate in continuous time rather than discrete time. *τ′* determines how rapidly firing rate approaches steady state for a constant synaptic current, and can be related to the membrane time constant (19). Although the relationship is not strict, we shall call *τ′* the membrane time constant. This is equivalent to convolving the instantaneous output of function F() with an exponential decay impulse response.

Equation 7 can be discretized by the Euler method, where *Δt′* = 1 time-bin = 5 ms. Discretized time is expressed as *t* instead of *t′* and the discretized form of the firing rate equation is:

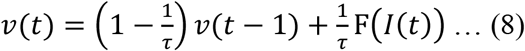

The dynamic network model incorporates units that behave like Equation 8. In recognition that the characteristics of units in our model may not exactly correspond to biophysical properties of neurons described by the firing rate equation, and to be consistent with the NRF model, we rename F() as g() and the current *I*(*t*) as activation *a*(*t*).

As a result, activation *a_j_*(*t*) of the *j*-th HU of the DNet model is the same as the NRF model,

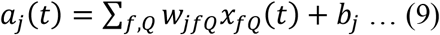

However, the output *v_j_*(*t*) is different from that of the NRF model, incorporating the discretized firing-rate equation.

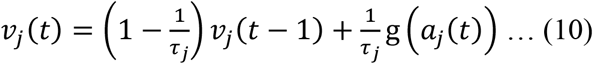

Similarly, the activation *a_o_*(*t*) of the OU is,

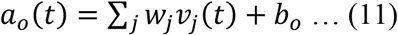

and the output v_*o*_(*t*) of the OU is,

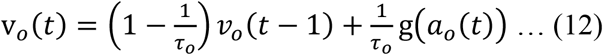

Here, *τ_j_* and *τ_o_* are the membrane time constants of *j*-th hidden unit and output unit respectively.

### The sDNet model

The firing-rate equation (Equation 7) assumes that the time constant that governs the relationship between the current and the firing rate is substantially larger than the time constant that governs the decay of post-synaptic potentials. However, if we assume the reverse, i.e., that the synaptic time constant is substantially larger, a different simple model of the relationship between the firing rate and the current becomes appropriate (19). This model is given by:

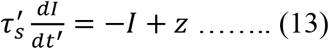

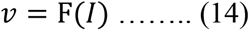

Here, *I* is synaptic current and *v* is post-synaptic firing rate, z is the immediate influence of the presynaptic spikes on the synaptic current, F() is the nonlinear function of synaptic current to post-synaptic firing rate, 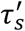 is synaptic time constant, and *t′* is time.

Similar discretization, like that used for the DNet model (see above), can be used for Equation 13 to use this for the units of a neural network model. Again, replacing the synaptic current, *I*, with the activation a, the activation of hidden units of the sDNet model can be modelled as the following:

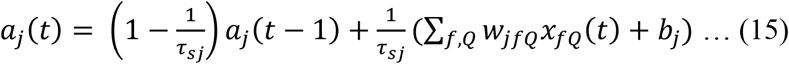

And the hidden unit output, replacing F() with g(), is given by:

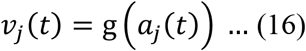

Hence, in the sDNet model, the activation of each HU, rather than the output of the nonlinearity, is convolved with an exponential decay impulse response.

Similarly, for the output units,

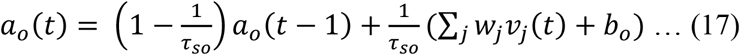

And the output,

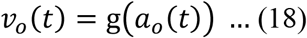

Here, τ*_sj_* and τ*_so_* are the synaptic time constants of *j*-th hidden unit and output unit respectively.

### Objective function of the NRF and DNet models

The free parameters *w_jfq_*, *w_j_*, *b_j_*, and *b_o_* (also τ*_j_* and τ*_0_* for the DNet and τ*_sj_* and τ*_so_* for the sDNet) were optimized by minimizing the squared error between *v_o_(t)* and *y(t)* subject to L1-regularization of the weights. Thus, the objective function is given by,

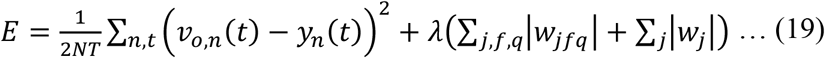

Here, *n* is included to indicate clip number, but was left out of other equations for simplicity. *N* is the number of clips used in training.

### Parameter estimation

All models except for the LN model are fitted by minimizing the objective function with respect to the free parameters using the sum-of-function optimizer algorithm (24). In using this algorithm, we take one clip to be one minibatch. The optimization algorithm requires calculation of the error gradients in respect to each of the parameters. For the NRF model, error gradients are calculated using standard chain rule. For the dynamic models, the process is similar (see below).

For the network models, before training, the weights were initialized by modified Glorot initialization from a uniform distribution ranging from 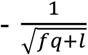 to 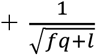, where *fq* is the number of input weights to a HU and *l* = 1 is the number of output weights from a HU. The biases were initialized similarly (25).

For the DNet model, to prevent the 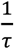 term running into mathematical error (when τ = 0), we substitute 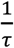 with 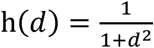. Parameters, d*_j_* and d*_0_* (for hidden and output units) were initialized from the square root of an exponential distribution with a mean of 1. Similar, substitution is used for the sDNet model.

### Error-gradient calculation for the DNet model

To estimate the parameters, the gradients of the error with respect to various parameters are calculated using a method similar to the one used for real time recurrent learning (26).

Substituting 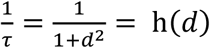, the dynamic equations for HUs (Equation 10) and OU (Equation 12) of the DNet model become respectively:

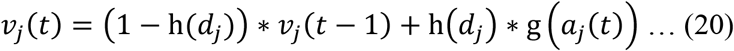

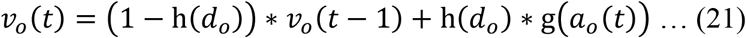

Using the chain rule, the gradient of the error term, E(*t*) at time *t* in respect to the weights of the output layer, *w_j_*,

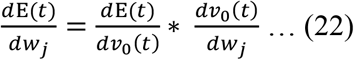

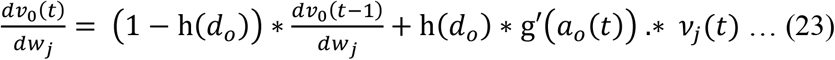

Similarly, the gradient of the error term, E(*t*) in respect to the bias of the output layer, *b_o_*,

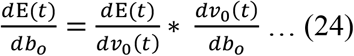

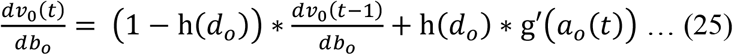

Finally, the gradient of the error term, E(*t*) with respect to the parameter, *d_o_*,

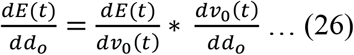

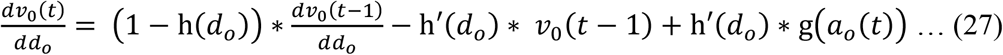

Hence, the gradient is passed forward from the current time step to the next time step to be used in calculation of the gradient at that new time step. At *t* = 1, the values of 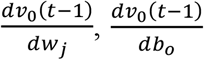 and 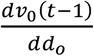 are undetermined. So, at time *t* = 1, the values of these terms are set to zero.

The gradients of the error term, E(*t*), with respect to the rest of the parameters of the DNet model and the sDNet model are obtained similarly.

### Cross-validation and model testing

From the data for the 20 sound stimuli, 4 were chosen as a test set which was not used during training and cross-validation. The cross-validation set (the remaining 16 stimuli) was used to fit the models using k-fold cross validation, where k = 8. The cross-validation set was randomly divided into a training set of 14 stimuli and a validation set of 2 stimuli. The model was trained on the training set for 18 different values of the hyperparameter λ. A log spaced range of lambda values was used, but with a somewhat lower density at the extremes. For the LN model, the exact values of λ used were: 1.00 × 10^−1^, 2.00 × 10^−2^, 1.17 × 10^−2^, 6.84 × 10^−3^, 4.00 × 10^−3^, 2.34 × 10^−3^, 1.37 × 10^−3^, 8.00 × 10^−4^, 4.68 × 10^−4^, 2.74 × 10^−4^, 1.60 × 10^−4^, 9.36 × 10^−5^, 5.41 × 10^−5^, 3.20 × 10^−5^, 6.40 × 10^−6^, 1.28 × 10^−6^, 2.56 × 10^−7^, and 5.12 × 10^−8^. For the rest of the models, the values of λ used were: 1.00 × 10^−3^, 2.00 × 10^−4^, 1.17 × 10^−4^, 6.84 × 10^−5^, 4.00 × 10^−5^, 2.34 × 10^−5^, 1.37 × 10^−5^, 8.00 × 10^−6^, 4.68 × 10^−6^, 2.74 × 10^−6^, 1.60 × 10^−6^, 9.36 × 10^−7^, 5.41 × 10^−7^, 3.20 × 10^−7^, 6.40 × 10^−8^, 1.28 × 10^−8^, 2.56 × 10^−9^, and 5.12 × 10^−10^. For each of the fitted models, neural responses were then predicted for the validation set, and the correlation coefficient between the actual neural responses and the prediction was measured. This process was repeated 8 times for different non-overlapping validation sets. The model was then retrained with the whole crossvalidation set using the λ value that provided the highest mean correlation coefficient over all 8 folds. Next, the retrained network was used to predict the neural responses to the test set. All the correlation coefficients and normalized correlation coefficients shown are for this held out test set, and all the model parameters shown are for the retrained network.

To ensure that all models receive the same amount of training data, models with different latency spans of STRFs are provided with exactly the same durations of neural response. This was done by clipping the necessary number of time points from the beginning of the neural response data.

### Effective hidden units

The hidden layer of each of the networks contained 20 Hus, but because their weights were L1-regularized, they tended to develop substantive weights only if they explained aspects of the neural response that were not explained by any other units. As a result, many HUs developed weights close to zero, and thus the network models tended to have only a small number of effective HUs. The ‘effectiveness’ of each unit *j* was calculated (20) as the variance over time of the unit′s weighted output *w_0_v_j_(t)*. Any HU with variance greater than 5% of the sum of the variances of all 20 HUs is considered effective.

### IE score

The IE score (20) measures whether the degree to which a HU is excitatory or inhibitory, on a scale between −1 and 1. Most inhibitory is IE = −1, when all of the input weights are negative or zero, and the output weight is positive, or when all of the input weights are positive or zero and the output weight is negative. Most excitatory is IE = 1, when all of the input weights are positive or zero, and the output weight is positive, or when all of the input weights are negative or zero and the output weight is negative.

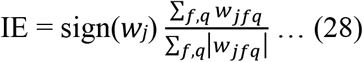

### Membrane time constant

Because the membrane time constant is modelled as 1 + *d*^2^ and the duration of each time step *Δt* = 5 ms, the value of membrane time constant in ms is 5 * (1 + *d*^2^).

### Knocking out large time constants of the DNet model

After fitting a network model, HUs with time constant greater than the length of the STRF are selected. The output weights connecting these units to the output unit are then set to zero. As a result, HUs with large time constants become inactive during the test process. Everything else in the model remained unchanged.

## RESULTS

The neural response dataset used in this study was recorded from the primary auditory cortical areas, A1 and AAF, of anesthetized ferrets in response to a diverse selection of natural sounds (20 stimuli of 5 s duration), including speech, animal vocalizations and environmental sounds (20,22). In total, 73 single-unit neural recordings showed sensitivity to the sound stimuli (see Methods), as measured by their noise ratios, and these were analysed in this study.

To probe the computations underlying stimulus integration, models were fitted to estimate the neural firing rate (averaged over stimulus repeats) of each neuron as a function of the preceding sound stimulation (Fig 1). First, for all models, the sound stimuli were pre-processed to generate a spectrogram-like representation (a cochleagram) that approximates the processing that occurs in the cochlea and auditory nerve. Second, using responses to 16 of the natural stimuli, a parameterized model was trained to estimate the firing rate as a function of the preceding cochleagram. After parameters of the models were fitted to the data, their fit quality was tested on a held-out test set composed of the responses to the remaining 4 natural stimuli.

**Fig 1:**
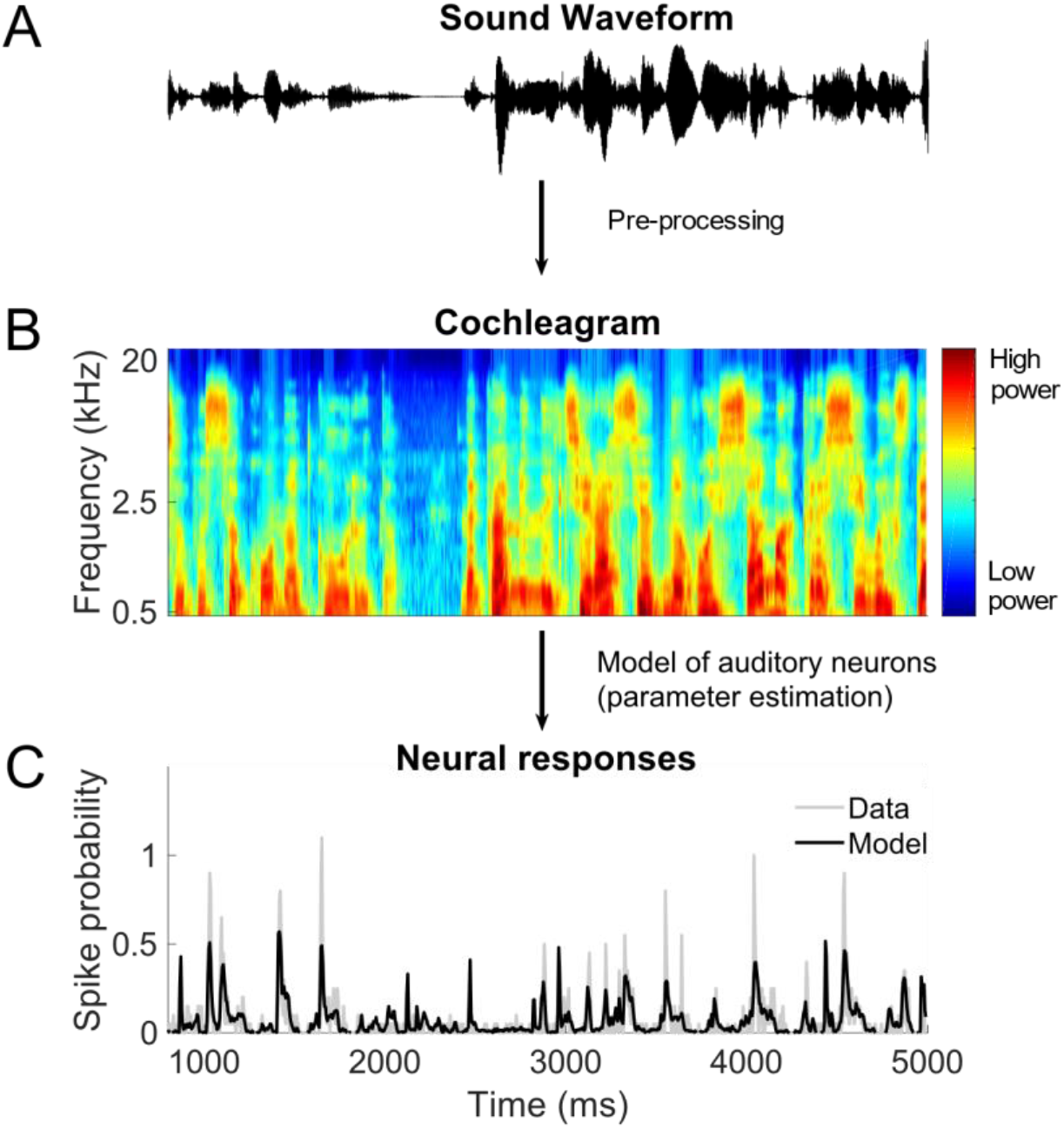
The models have two-stages. A. The first stage involves pre-processing that mimics the auditory periphery, where the sound waveform is converted into a time-frequency representation (cochleagram). B. The output of the first stage is then used as input to a model that is fitted to the neural responses of auditory cortical neurons. C. After parameter estimation, the model was used to predict neural responses to arbitrary stimuli.

### Using standard models, access to 200 ms of recent stimulus history is needed to achieve best prediction

For each neuron, an LN model was fitted to estimate its firing rate as a function of the preceding cochleagram. The LN model consists of an STRF, followed by a sigmoid nonlinearity (Fig 2A). As mentioned in the Methods, the STRF is the weighted sum of the cochleagram over frequency components at a span of latencies. The effect of different spans of latencies (lengths) of the STRF was investigated. For the held-out test set, and averaged over the 73 neurons, the normalized correlation coefficient, CC_norm_ (27,28), between the actual neural responses and the prediction of LN models with 25, 50, 100, 200, and 400 ms long STRFs are respectively 0.51, 0.64, 0.69, 0.71, and 0.71 (Fig 2B), where 1.0 is the maximum achievable prediction given the estimated neuronal noise and 0.0 indicates that there is no correlation between actual and predicted neural responses. The means of non-normalized Pearson correlation coefficients (CC) are 0.40, 0.50, 0.54, 0.55, and 0.55 respectively. Fig 2C shows STRFs of some example neurons estimated by the LN model.

**Fig 2:**
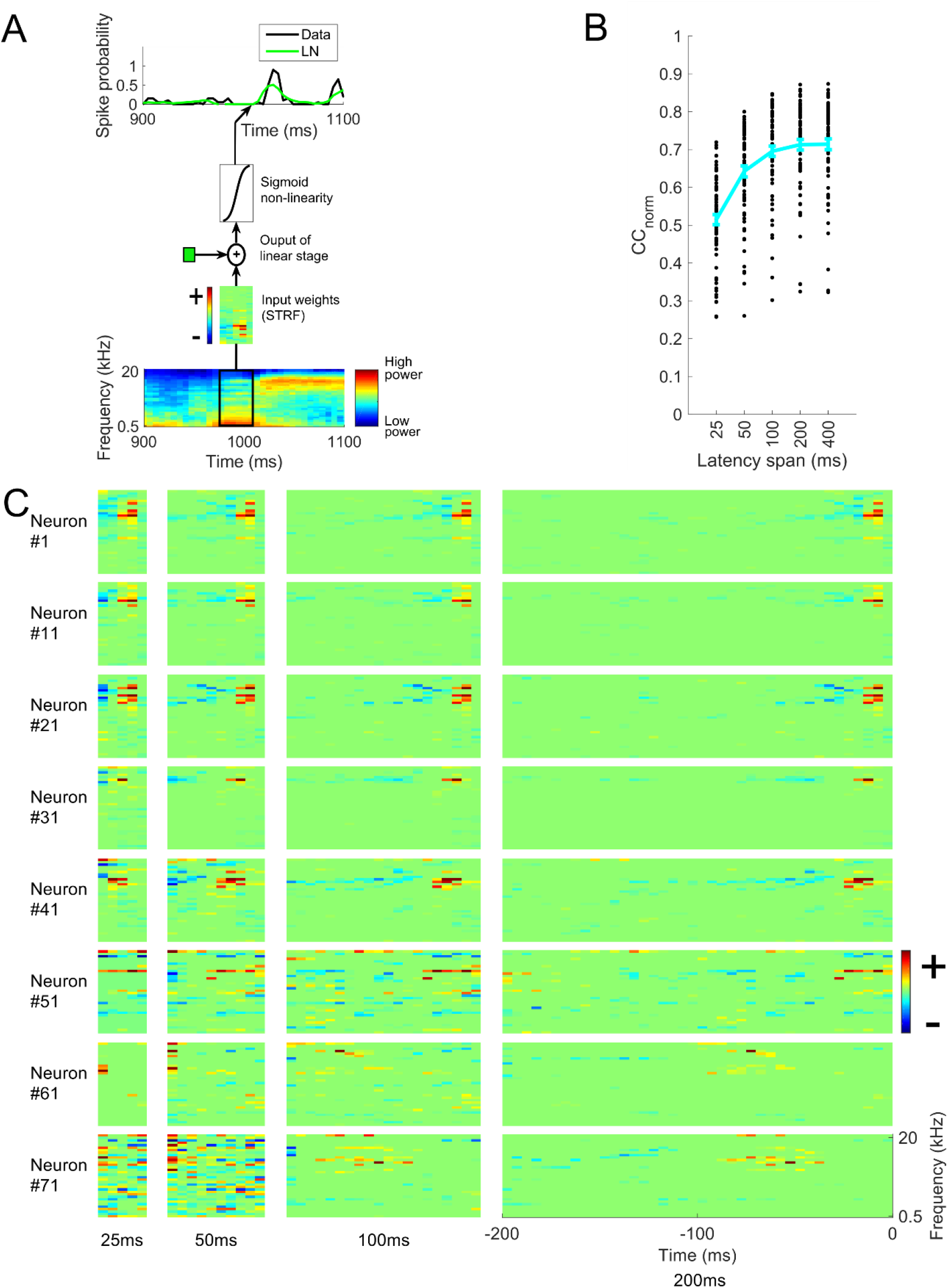
Prediction of neural responses by the linear non-linear (LN) model. A. The LN model, which consists of a spectrotemporal receptive field (STRF) followed by a sigmoid nonlinearity. B. Normalized correlation coefficients (CC_norm_) between neural responses and the prediction of the LN model with different latency spans. Each black dot represents a neuron, the curve indicates the mean over 73 neurons and error bars indicate standard error of the mean. C. STRFs of some example neurons, for LN models with different latency spans. Red, excitatory fields, blue, inhibitory. Excitatory and inhibitory fields extend up to 100 ms in most cases and up to 200 ms in some cases.

A network receptive field model (the NRF model) was also developed and fitted to each neuron. This model consisted of the weighted sum of multiple LN-like units (similar to 19). The NRF model is essentially an artificial neural network with a single hidden layer, using a sigmoid non-linearity in its hidden units and its single output unit (Fig 3A). The NRF model makes use of multiple component STRFs, which allows modelling of neuronal sensitivity to non-linear interactions of multiple features (20). STRFs of at least 400 ms are needed for best performance in predicting neural responses (Fig 3B). For the test set, the mean CC_norm_ between neural responses and the model predictions made by NRF models with 25, 50, 100, 200 and 400 ms long component STRFs are 0.51, 0.64, 0.70, 0.72 and 0.73, respectively. The mean CC are 0.40, 0.50, 0.54, 0.55 and 0.56. Fig 3C shows the input weights (analogous to STRFs) for the ‘effective’ HUs (see Methods) of the model-fit for one example unit, obtained using the NRF model.

**Fig 3:**
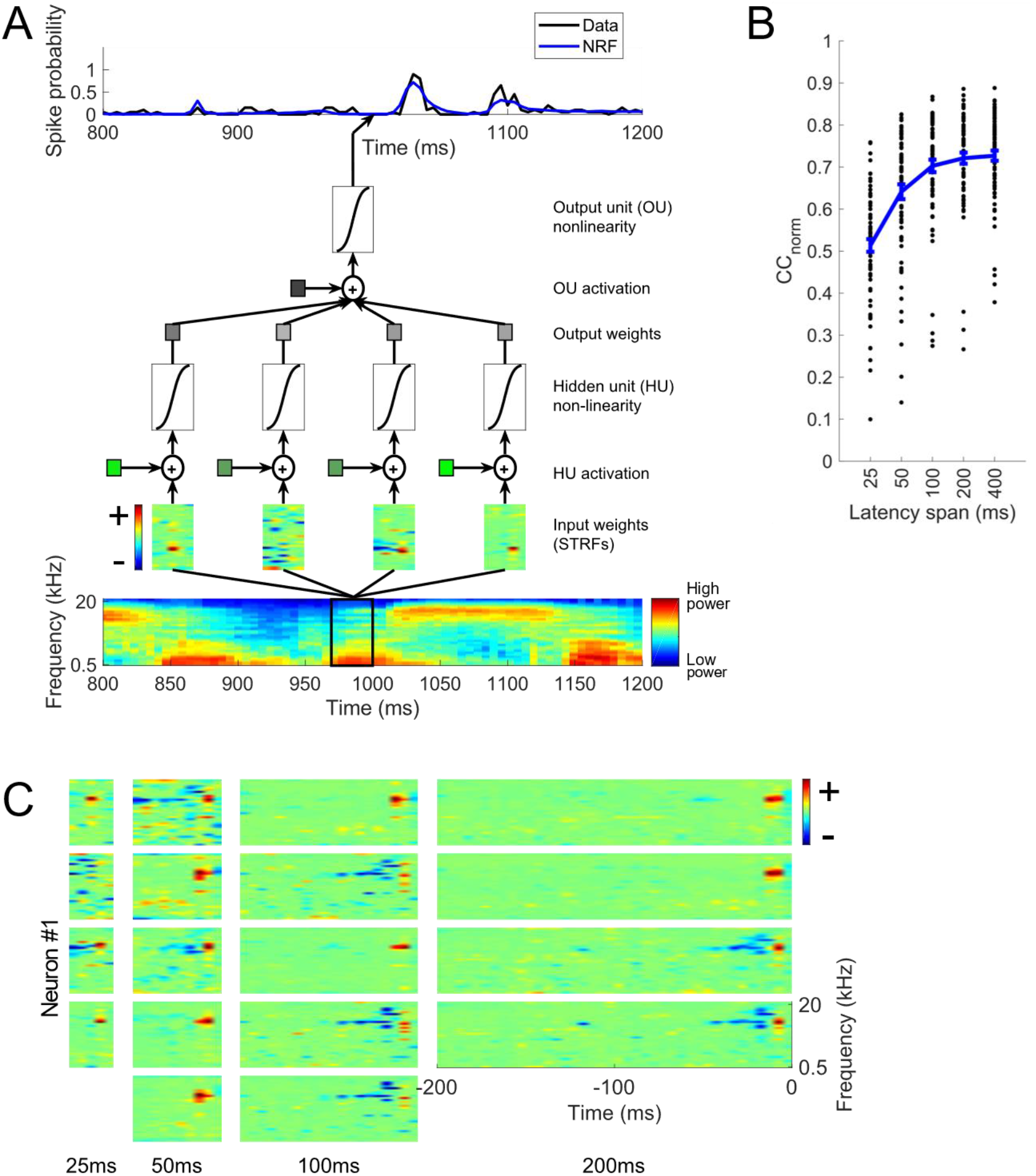
Prediction of neural responses by the network receptive field (NRF) model. A. The NRF model, which involves a weighted sum of multiple LN-model like units, each with an STRF. B. Normalized correlation coefficients (CC_norm_) between neural responses and the prediction of the NRF model with different latency spans. Each black dot represents a neuron, the curve indicates the mean over 73 neurons and error bars indicate standard error of the mean. C. Component STRFs of an example neuron. Each column represents an NRF model with different latency spans.

### The DNet model can characterize neural responses using very short duration STRFs

Although both the LN and NRF models can capture dependence on stimulus history using a span of input latencies extending up to at least 200 ms, there is little biological evidence for such a wide span of input latencies (see Introduction). With the aim of better understanding the biological underpinnings of stimulus history dependence in auditory cortical responses, a model with a dynamic aspect was developed that integrates the output of its component units over time in a manner similar to those of real neurons (see Methods). Each unit in the DNet model includes exponentially-decaying response integration, motivated by the integrative properties of real neuronal membranes (Fig 4A). This model is in essence the NRF model but with dynamic units, each modelled by the dynamic firing-rate differential equation (Equation 7) (19). That is, each unit of the model applies an exponentially decaying impulse response to the output of their non-linear activation function. Each unit’s decay rate is governed by its individually fitted time constant, *τ* (see Methods).

**Fig 4:**
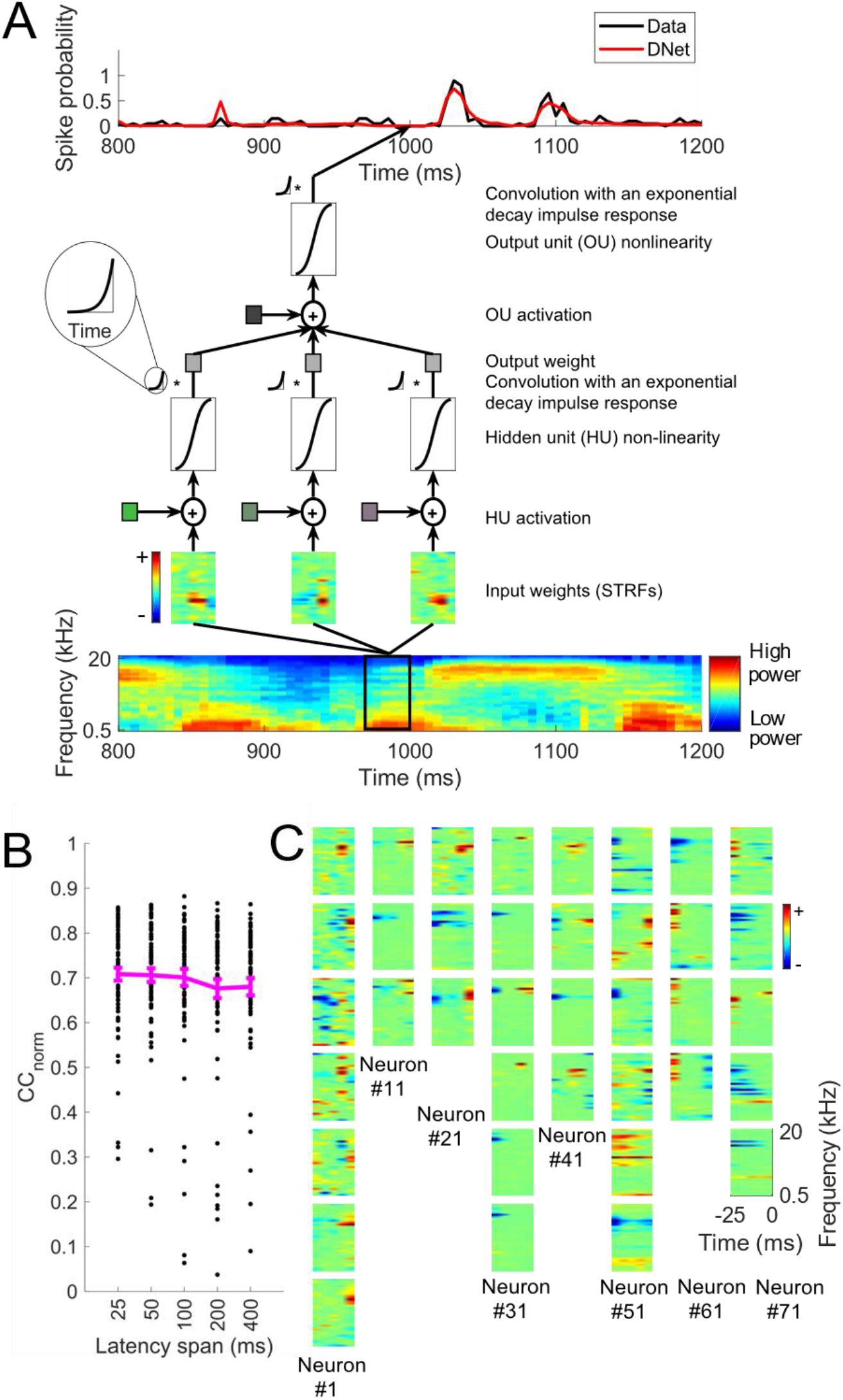
Prediction of neural responses using a dynamic network (DNet) model. A. The DNet model. The cochleagram is passed through a set of linear-nonlinear filters, whose response is then integrated over time using an exponentially-decaying impulse response to produce the hidden units’ outputs. A weighted sum of these outputs is passed through a similar output unit to make a prediction of the neural response. B. CC_norm_ for the DNet model with different latency spans. Each black dot is a neuron, the curve indicates the mean over 73 neurons and error bars are standard error of the mean. C. The input weights (STRFs) for the ‘effective’ hidden units (see Methods) of 8 example neurons (each column is a different neuron) for the DNet model.

We found that the mean CC_norm_ between the actual neural responses and the predictions of the DNet model with 25, 50, 100, 200 and 400 ms STRFs are respectively 0.71, 0.71, 0.70, 0.68 and 0.68 (Fig 4B). The mean CC are 0.55, 0.55, 0.54, 0.52 and 0.52. Because the 25-ms model performs the best and the biological range of latencies are typically less than 30 ms (Bizley et al., 2005), we focused on the 25-ms DNet model for further analysis. Fig 4C shows the input weights (analogous to STRFs) for the ‘effective’ hidden units (see Methods) of the model fits for 8 example neurons, using a DNet model with 25-ms long STRFs.

The 25-ms DNet model outperforms the DNet with longer (200 ms and 400 ms) STRFs, as measured by prediction of neural responses on a held-out test set averaged over all 73 neurons (Fig 5A). It also achieves performance similar to the best performing LN model, but performs a little worse than the best performing NRF model (Fig 5A). This can be seen qualitatively in the time course of the prediction for an example neuron and stimulus in Fig 5B. A neuron by neuron comparison shows that this result is consistent for most neurons (Fig 5C and 5D). Also, the 25-ms LN and NRF models are outperformed by a large margin by the 25-ms DNet model (Fig 5A). A neuron by neuron comparison shows that this result is also consistent for almost all neurons (Fig 5E and 5F).

**Fig 5:**
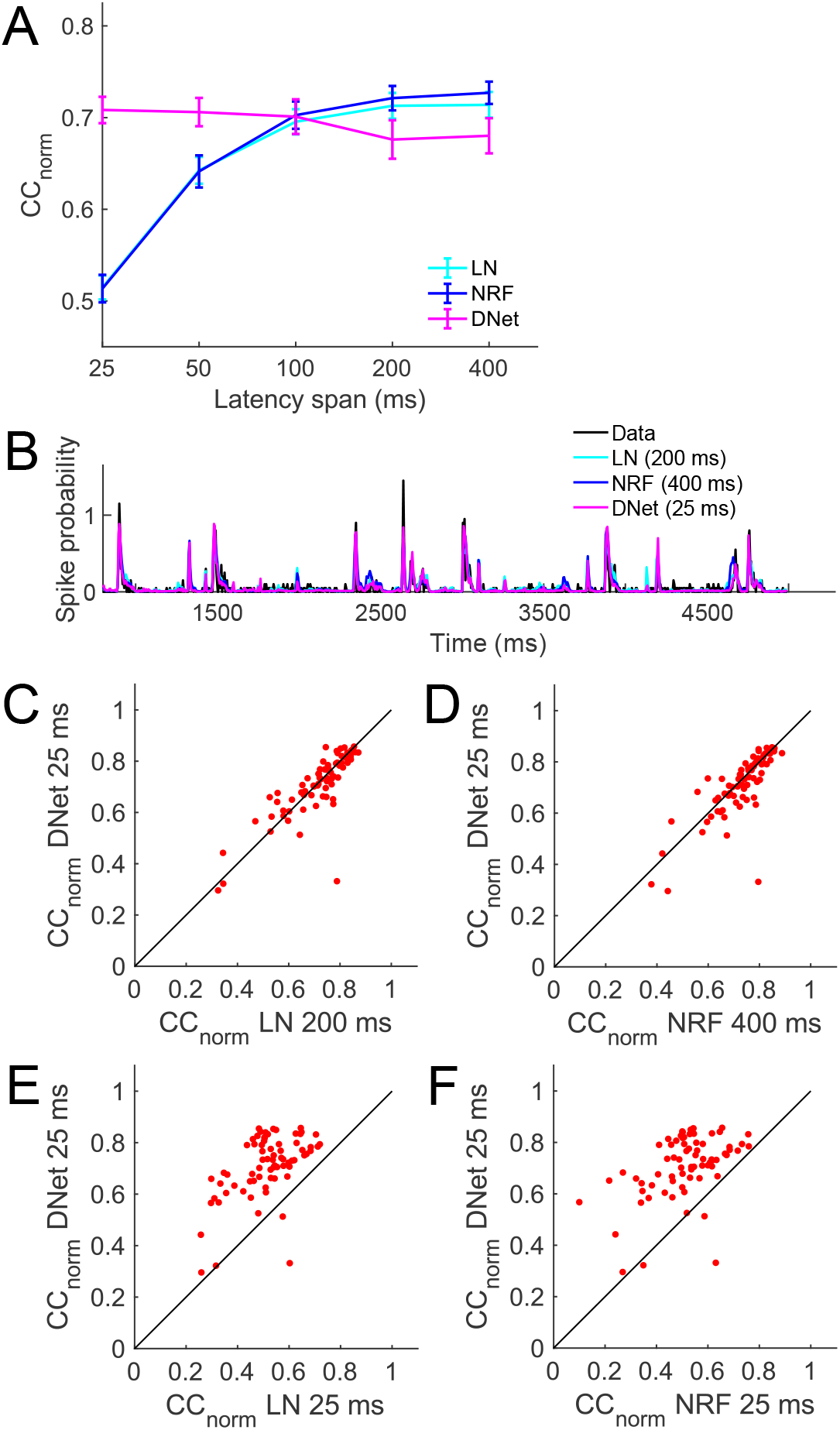
Comparison of the performance of the DNet model with other models. A. Mean CC_norm_ of linear-nonlinear (LN), network receptive field (NRF) and dynamic network (DNet) models. B. Time course of predictions by best performing versions of each of the LN, NRF and DNet models, compared with real neural responses for an example neuron and stimulus. C. Neuron by neuron comparison of CC_norm_ values between the 25-ms DNet model and 200-ms LN model. Each red dot is a neuron. D. Neuron by neuron comparison of CC_norm_ values between the 25-ms DNet model and 400-ms NRF model. E. Neuron by neuron comparison of CC_norm_ values between the 25-ms DNet model and 25-ms LN model. F. Neuron by neuron comparison of CC_norm_ values between the 25-ms DNet model and 25-ms NRF model.

### Network structure and response integration are both required for an effective DNet

To understand how much of an effect HUs of the DNet model with long time constants have on the model’s predictions, the fitted model was modified by ‘knocking out’ (see Methods) HUs with time constants longer than the duration of their STRFs. This was achieved by setting the output weights of those units to zero, while keeping all other parameters intact. We found that the 25-ms DNet model is the DNet whose CC_norm_ is most reduced by this alteration (Fig 6A).

**Fig 6:**
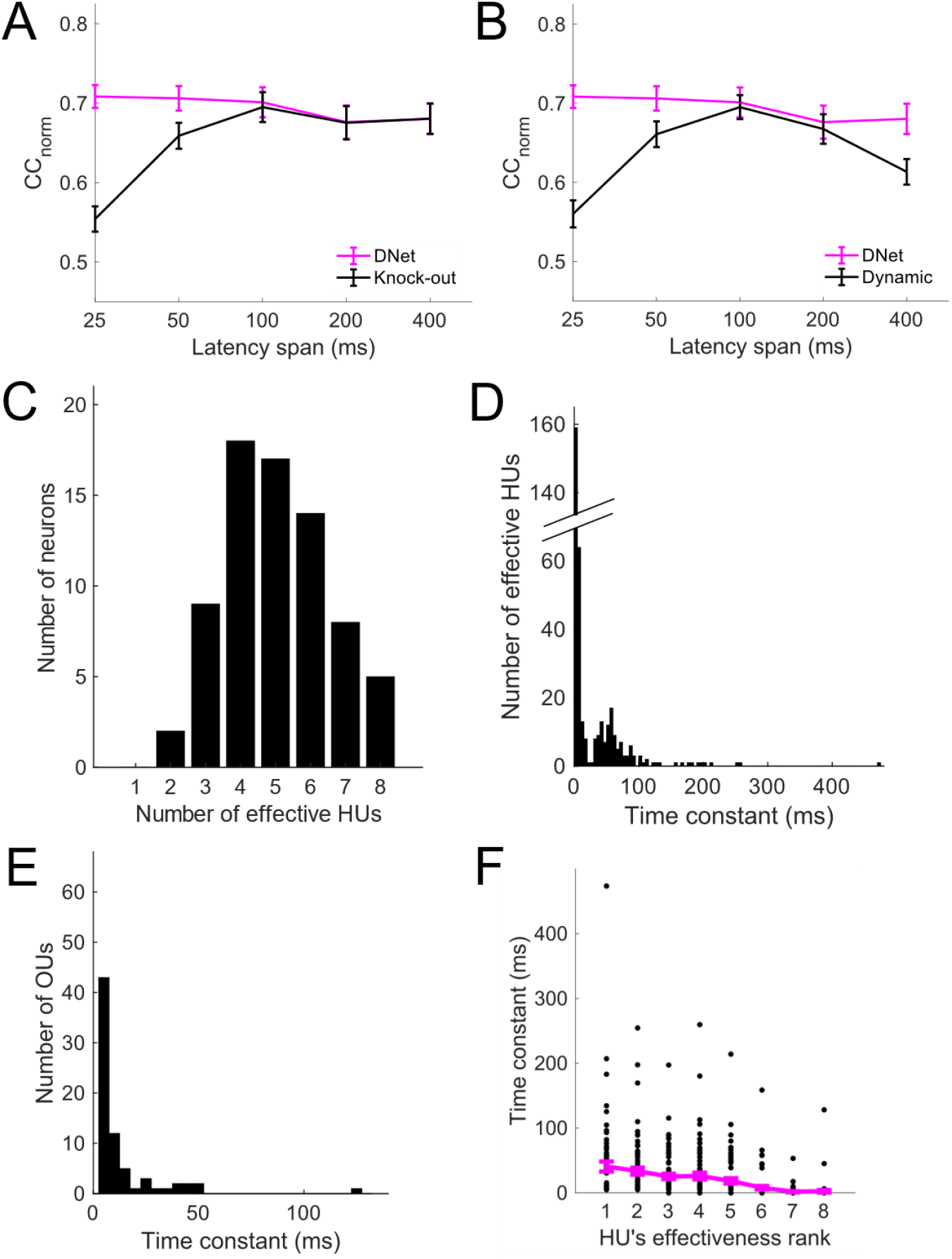
Multiple time constants in a network architecture are required to capture the dependence of neural response on stimulus history. A. DNet model performance when large time constants are knocked out. B. Performance of the single-unit dynamic model. C. Number of effective hidden units (HUs) used to model each of the 73 neurons. D. Distribution of time constants of HUs. E. Distribution of time constants of the output units (OUs). F. Relationship between time constant and HU effectiveness.

A single-unit dynamic model was also developed to examine the effect of a time constant in an LN-like single STRF model. Prediction performance of this model was found to be worse than the LN model (Fig 6B). This indicates that both long time constants and a network architecture are needed for the 25-ms DNet model to achieve prediction performance similar to that of standard models.

Although the DNet model had 20 hidden units, the L1 regularization eliminated unnecessary units. Hence there were fewer effective HUs, and for the 25-ms DNet model this number was found to be between 2 and 8 (Fig 6C). We defined an effective HU as one whose effectiveness (variance of weighted output, see Methods) exceeded a certain threshold.

In the 25-ms DNet model, the time constants of the effective HUs ranged between 5 and 475 ms, often far greater than the duration of the STRFs (Fig 6D). The time constants of the OUs of the DNet model ranged from 5 to 130 ms (Fig 6E). To investigate whether long time constants are associated with effective HUs, for each neuron HUs were ranked in order of effectiveness and the mean of the time constants of all HUs in the same rank (over all neurons) were calculated. This showed that high-ranked HUs tend to have long time constants (Fig 6F).

### Hidden units with long time constants tend to be more inhibitory

We identified the effective HUs with long time constants, i.e., where the value of time constant is greater than the length of the STRF (25 ms). To test whether a relationship exists between the time constant and unit’s inhibitory/excitatory nature, we classified the effective HUs of the 25-ms DNet model into excitatory or inhibitory using an IE score (see Methods). If the output weight is positive, an IE score of 1 means a unit with entirely excitatory influence and an IE score of −1 means a unit with entirely inhibitory influence. We found that almost all units having large time constants are inhibitory in nature (Fig 7).

**Fig 7:**
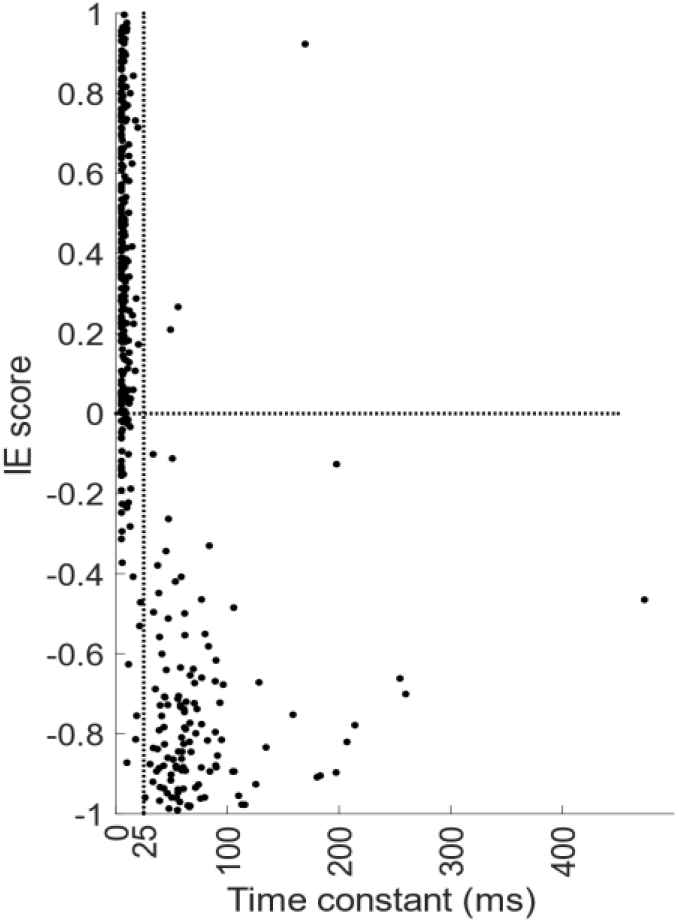
Relationship between inhibitory/excitatory (IE) score and time constant. Almost all the HUs with time constants greater than the duration of the STRFs (units right of the vertical dotted line, the duration of the 25 ms STRFs) are inhibitory in nature. IE score = inhibitory/excitatory score.

### Membrane time constant versus synaptic time constant

In the DNet model, the integration governed by the time constant occurs after each unit’s nonlinearity. Although the time constant is conceived and implemented in a manner consistent with a membrane constant, within the network it may capture other dynamic neural phenomena such as synaptic time constants, forms of a channel-based or synaptic adaptation, or certain forms of network recurrency. An alternative network model places the integration just before the nonlinearity, which is more consistent with the time constant being synaptic in nature (19). To investigate how much of the neural activity can be explained by such a model, we modified the DNet model to form the sDNet model (synaptic DNet model), with each unit governed by Equations 13 and 14 (19).

The sDNet model achieves best performance in predicting neural responses when the duration of the STRFs is 200 ms (CC_norm_ = 0.71) (Fig 8A). Although the sDNet with 25-ms STRFs predicts neural responses better (CC_norm_ = 0.67) than the 25-ms LN model (CC_norm_ = 0.51) and the 25-ms NRF model (CC_norm_ 0.51), it is worse than the 25-ms DNet model (CC_norm_ 0.71). This suggests that a large percentage of the variance of neural responses can be explained by a time constant before non-linearity, but that having a time constant after the non-linearity further improves prediction.

**Fig 8:**
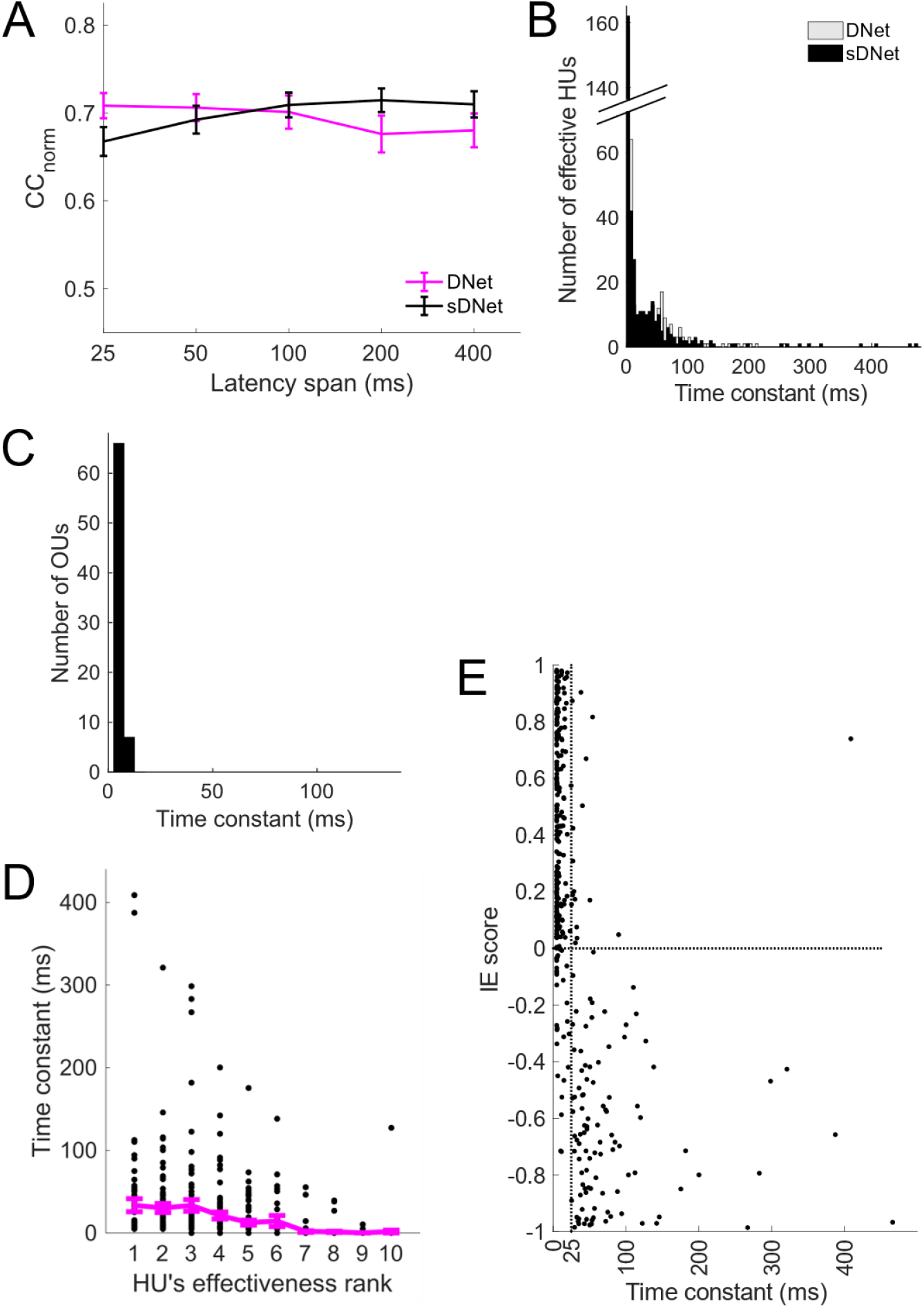
A modified DNet model with synaptic time constants (sDNet model). A. Performance of the sDNet model. B. Distribution of the values of time constants of hidden units (HUs) in the sDNet model (black bars). Gray bars in the background represent time constants of HUs in the DNet model. C. Distribution of the values of time constants of the output units (OUs). D. Relationship between a HU’s time constant and its effectiveness. E. Relationship between a HU’s inhibitory/excitatory (IE) score and its time constant. Most of the HUs with time constants greater than the duration of the STRFs (units right of the vertical dotted line) are inhibitory in nature.

While the largest time constants of the effective HUs of the sDNet model can be as large as ~470 ms (Fig 8B), the distribution of time constants of output units have smaller values compared to the DNet model (Fig 8C). As with the DNet model the most effective hidden units are those with longer time constants (Fig 8D) and units with long time constants are inhibitory in nature (Fig 8E).

## DISCUSSION

We find that a model of primary auditory cortical neurons which incorporates exponentially-decaying memory processes – the DNet model – can capture temporal aspects of the receptive field using very short (25 ms) component STRFs in a network. Standard models such as the LN model (and the NRF model) model temporal receptive field of neurons using delay lines whose latencies extend up to a few hundred milliseconds. Our results suggest that, beyond 25ms, the temporal receptive field can be modelled more succinctly by exponentially-decaying memory processes than by delay lines.

When auditory neurons are fitted using a standard LN model, the resulting STRFs often contain narrowband inhibitory fields that decay over approximately 200 ms. In contrast, we find that the STRFs of the DNet model have short inhibitory fields with long time constants. This suggests that these temporally extended inhibitory fields in STRFs reflect neural mechanisms that can be approximated by exponentially-decaying memory processes.

There are many biological mechanisms which may underlie these exponentially-decaying memory processes. Neurons have various dynamic integrative and adaptive characteristics originating from membrane dynamics, short-term synaptic plasticity, post-synaptic potentials, channel-based adaptation and recurrent network effects (19). In a previous study, a model with synaptic depression has been shown to partially capture the integration of stimulus history in A1 (29). Here, we found that a better prediction of neural responses can be achieved by a model based on the integrative properties of neural membranes (the DNet model) than by the one based on the dynamics of synaptic potentials (the sDNet model). We note, however, that although the DNet model was developed with membrane dynamics in mind, aspects of the above listed dynamic processes could be captured by the same exponentially-decaying memory.

The firing rate of A1 neurons has been demonstrated to depend on stimulus history up to ~4 s in the past (17). This dependence cannot be fully captured by linear models such as STRFs (17). Although the DNet model can capture more succinctly aspects of neural dependence on stimulus history, this indicates that there are some nonlinear dynamic processes that provide history dependence which cannot be captured either by linear or LN models, or by the DNet model. Perhaps this is because the DNet model has limited capacity to capture temporal dynamics that are not exponential in nature.

Much of the relationship between stimulus and neural responses results from neurons being embedded in a network structure and from the nonlinear properties of the units in this network. From the auditory periphery to the A1, there are numerous feedforward, feedback and lateral connections. Many previous models of auditory cortical neurons have not explicitly represented this network character. However, some recent models have incorporated non-linear contextual effects (14,22,30–32). Furthermore, a few models do explicitly incorporate multiple receptive fields (33–39) or more extensive network structure (20), which perform better in predicting neural responses of A1/AAF neurons than STRF models (20,33).

When modelling the dependence of neural responses on stimulus history, one approach is to include the history of a neuron’s spikes and those of its recorded neighbours in a generalized linear model (GLM) model (31,40). However, unlike the DNet model, this approach does not model multi-layer network structure with hidden units that are inferred but not recorded. Modelling hidden network structure has been applied to the salamander retina, where in vitro responses to simple artificial stimuli are well predicted by a network model (LNFDSNF) with various non-linearities and response delays mimicking the retinal pathway (41). The LNFDSNF model is particularly relevant to our study because it is a network model whose units have some memory of their past activity, though the form of this memory differs from that of the DNet model. The LNFDSNF model units have a feedback kernel that acts before the nonlinearity - in contrast, the DNet model units integrate their activity after the nonlinearity using an infinite impulse response that decays exponentially.

The succinct and biologically-motivated description of neural responses provided by the DNet model provides a basis from which to develop further models to account for the ~30% of the CC_norm_ that remains to be explained. One variant of the model that can be developed involves a network with multiple hidden layers where the units each have just a single latency (i.e. each unit in the first hidden layer uses just a spectral receptive field at a single delay). This approach provides an opportunity to build deeper neural-network-like models with fewer parameters, potentially overcoming overfitting problems for deep network models of neural responses (42).

Neurons are intrinsically dynamic systems, each with memory of past events. Furthermore, neurons are embedded in networks. We show that models incorporating both these characteristics can be very valuable for explaining moment to moment *in vivo* responses of neurons in the auditory cortex. This is a more biologically plausible way of capturing the history dependence of neural responses than a set of delay lines. In artificial neural networks, memory is usually implemented through recurrent connections between units. However, real biological neurons also have intrinsic memory processes with particular characteristics. Including such single-unit memory has advantages for explaining the responses of biological neurons and may also be useful in artificial neural networks.

## Author contributions

N. S. H originated project; M. R. and N.S.H. designed research; M.R., B.D.B.W. and N.S.H. performed research; M.R., B.D.B.W. and N.S.H. contributed unpublished reagents/analytic tools; M.R. and N.S.H. analyzed data; M. R., B.D.B.W., N.S.H., A.J.K. wrote the paper.

## Acknowledgements

B. D. B. W., N. S. H., and A. J. K were supported by Wellcome Trust funding (WT108369/Z/2015/Z). M.R. was supported by the Clarendon Fund Scholarship.

